# Disruption of Human *astn2* Function by ZIKV *ns4b* Gene as a Molecular Basis for Zika Viral Microcephaly

**DOI:** 10.1101/054486

**Authors:** Bhaskar Ganguly, Enakshi Ganguly

**Author notes:** Corresponding author, address for correspondence: C/o Late *Sri* Bisweswar Gangopadhyay, D-04, Alliance Kingston Estate, Rudrapur 263153. Uttarakhand. INDIA.

## Abstract

The present Zika virus (ZIKV) pandemic is being associated with increased incidence of microcephaly in newborns. However, a molecular basis for such pathogenesis is distinctly lacking. Comparative nucleic acid sequence analysis showed similarity between regions of non-structural protein 4B (*ns4b*) gene of ZIKV and human astrotactin2 (*astn2*) gene. Based on these findings, a molecular target of Zika viral microcephaly is being proposed.

## 1. INTRODUCTION

Given the present Zika virus (ZIKV) pandemic, there is an accentuated interest in its pathobiology. ZIKV isa mosquito-borne Flavivirus, whose natural transmission cycle involves mainly vectors from the Aedes genus and monkeys. The ZIKV genome consists of a singlestranded positive sense RNA molecule, about 11 Mb in size, with two flanking non-coding regions (5′ and 3′ NCR) and a single long open reading frame encoding a polyprotein: 5′-C-prM-E-NS1-NS2A-NS2B-NS3-NS4A-NS4B-NS5-3′, that is cleaved into capsid (C), precursor of membrane (prM), envelope (E) and seven non-structural proteins (NS 1 through 5). Humans were considered occasional hosts of the virus and till 2007 only 14 human cases had been reported. The potential of ZIKV as an emerging human pathogen was first realized during the year 2007 epidemic at Yap Island in Micronesia involving 49 confirmed cases of morbidity and 73% seropositivity in the adult population.^1^ Before April, 2015, no case had been reported in Brazil. However, between April and November, 2015, 18 of the 27 Brazilian states reported ZIKV. The most arresting of all the present concerns regarding the manifestation of ZIKV is its association with microcephaly in newborns. After ZIKV emerged in Brazil, a 20-fold annual increase of microcephaly cases was observed.^2^ A relationship between ZIKV and microcephaly has been confirmed, and WHO has issued an epidemiological alert about the association of ZIKV infection with microcephaly,^3^ although a molecular basis for associating the condition with the pathogen is still lacking.

Pathogens and hosts have evolved in each other’s presence and parsimonious analysis of the two by nucleic acid sequence comparison can uncover hitherto unknown interaction networks.^4^ Therefore,it was posited that a parsimonious association between ZIKV and humans may lead to a patho-molecular basis for microcephaly.

## 2. METHODS

The ZIKV genome sequence [RefSeq: NC_012532.1]^5^ was used to query the human non-redundant nucleotide sequence database using blastn^6^ for finding ‘somewhat similar sequences’ keeping all search parameters at their default value. Sequences in the human genome that showed alignment with ZIKV genome sequences were manually annotated using extensive literature search for finding a possible role in pathogenesis of microcephaly.

## 3. RESULTS AND DISCUSSION

BLAST search revealed sequence matches of different parts of ZIKV genome with several different targets in the human genome. One of these matches of functional significance in the etiopathogenesis of microcephaly was that of ZIKV *ns4b* gene with *astrotactin 2 (astn2)* gene(Figure 1).

**Figure 1.**
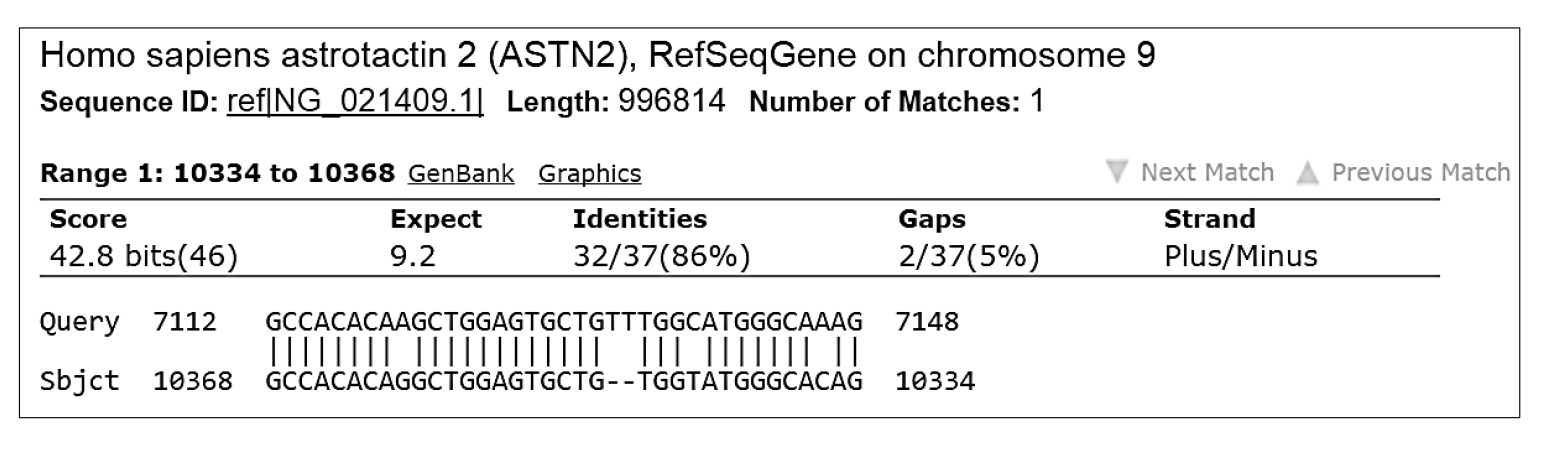
Blastn output showing local alignment of ZIKV *ns4b* (query)with human *astn2*(subject)

An expect value of 9.2 for the alignment appears high but is likely due to the large disparity in the size of the query and the alignment with the subject. When a reciprocal blastn search was performed using the *astn2* gene sequence to query the ZIKV non-redundant sequences, the same alignment was achieved with an expect value of 0.049 (data not shown). The non-structural protein 4B (NS4B) protein is small and poorly conserved among the Flaviviruses. NS4B contains multiple hydrophobic potential membrane spanning regions. NS4B may form membrane components of the viral replication complex and could be involved in membrane localization of NS3.^7^ ASTN2 is a protein that is expressed at high levels in migrating, cerebellar granule neurons during developmental stages when glial-guided migration is ongoing and mutations of *astn2* gene are associated with microcephaly in humans.^8^

Based on the above finding, it first appears that both ZIKV NS4B and host ASTN2, being transmembrane proteins, may co-localize and interact with each other in such a manner as to annul the physiological role of ASTN2 but a closer analysis of the alignment reveals that the part of*astn2* gene exhibiting sequence identity with ZIKV *ns4b* is located within a 123 kB intron (intron 1) coding region rather than forming a part of the exon coding region (Figure 2). Thus, it can be predicted that ZIKV *ns4b* alters *astn2* function through pre-translational interactions to cause microcephaly. The exceptional ability of ZIKV as a Flavivirus to exist within the host cell nucleus strengthens this model of pathogenesis.^9^ However, it is very difficult to speculate on the nature of the exact pre-translational interaction between the two genes that results in disruption of *astn2* function.

**Figure 2.**
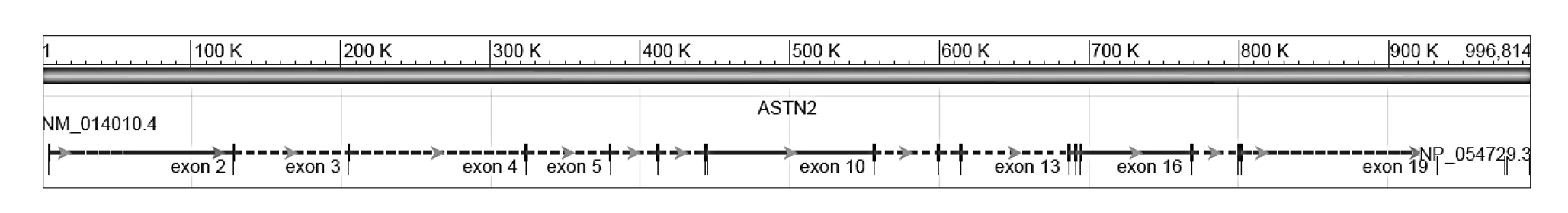
The gene structure of human *astn2*; the gene comprises of 22 exons. ZIKV *ns4b* is likely to interact with the first intron, about 123 kB in size, close to the first exon at about 10.3 K on the scale shown.

The 123 kB intron commonly features in many proteins of the epidermal growth factor receptor(EGFR) family,^10^ and the alignment site within the intron is unusual in having a high GC content (22/37=59%); higher GC content is typically associated with shorter introns.^11^ The presence of long non-coding, anti-sense RNA gene, PAPPA-AS1, which is differentially expressed in placenta and brain, juxtaposed with *astn2* on human chromosome number ^9^,^12^ may also bear excitating implications in the molecular pathogenesis of ZIKV microcephaly. Biological studies abounding the determination of the copy numbers of functional *astn2* mRNA and ASTN2 protein in developing fetuses that are at risk of developing microcephaly due to maternal ZIKV exposure would be required to confirm this model of pathogenesis. If the proposed model of pathogenesis for zika viral microcephaly involving *astn2* and ZIKV *ns4b* is confirmed, it would allow development of new medical interventions for complementation of *astn2* function towards successful prevention of ZIKV microcephaly.

